# Control of Systemic Iron Homeostasis by the 3’ Iron-Responsive Element of Divalent Metal Transporter 1 in Mice

**DOI:** 10.1101/2020.02.20.957779

**Authors:** Elisabeth Tybl, Hiromi Gunshin, Sanjay Gupta, Tomasa Barrientos, Michael Bonadonna, Ferran Celma Nos, Gael Palais, Zoubida Karim, Mayka Sanchez, Nancy C. Andrews, Bruno Galy

## Abstract

Divalent metal transporter 1 (DMT1) is essential for dietary iron assimilation and erythroid iron acquisition. The 3’ untranslated region of the murine DMT1 mRNA contains an iron responsive element (IRE) that is conserved in humans but whose functional role remains unclear. We generated and analyzed mice with targeted disruption of the DMT1 3’IRE. These animals display hypoferremia during the suckling period, associated with a reduction of DMT1 mRNA and protein in the intestine. In contrast, adult mice exhibit hyperferremia, accompanied by enlargement of hepatic and splenic iron stores. Intriguingly, disruption of the DMT1 3’IRE in adult animals augments intestinal DMT1 expression, in part due to increased mRNA translation. Hence, during postnatal growth, the DMT1 3’IRE promotes intestinal DMT1 expression and secures iron sufficiency; in adulthood, it suppresses DMT1 and prevents systemic iron loading. This work demonstrates that the 3’IRE of DMT1 plays a role in the control of DMT1 expression and systemic iron homeostasis, and reveals an age-dependent switch in its activity.

**Key points:**

- Targeted mutagenesis of the 3’IRE of DMT1 in mice reveals its importance for maintenance of systemic iron homeostasis.
- The 3’IRE stimulates intestinal DMT1 expression and prevents hypoferremia during early life, but exerts opposite effects in adulthood

## Introduction

Tight control of intestinal iron absorption is required to avoid both iron insufficiency^1^ and excess^2^. Dietary non-heme iron is taken up by absorptive enterocytes via the apical iron transporter DMT1 (a.k.a. SLC11A2)^3-5^, and transferred into the circulation by ferroportin (FPN, a.k.a. SLC40A1)^6^, with the help of a ferroxidase, hephaestin (HEPH)^7^. FPN activity is controlled by the liver hormone hepcidin^8^, but DMT1 seems regulated locally via mechanisms operating within enterocytes^9,10^. DMT1 mRNA exists in four isoforms that differ in their 5’ and 3’ ends^11,12^. 3’ end diversity results from alternative usage of splicing and polyadenylation sites, and yields isoforms that either contain or lack a conserved iron responsive element (IRE) in their 3’ untranslated region (UTR). IRE-containing isoforms are predominant in duodenal enterocytes^12^. IREs are stem-loop structures that interact with iron regulatory proteins (IRPs, a.k.a. ACO1 and IREB2) in iron-depleted cells^13^. IRP binding to multiple IREs in the 3’-UTR of the transferrin receptor (TFRC) mRNA limits its degradation by Regnase-1 (a.k.a. ZC3H12A)^14^. The presence of an IRE-like motif in *DMT1* suggests that DMT1 could be regulated by IRPs, similar to TFRC. However, the single DMT1 IRE contains an additional 3’-bulge in its upper stem^15^, and DMT1 mRNA seems to lack a Regnase-1 binding site^16^. Importantly, DMT1 expression only responds to iron fluctuation in a subset of cell lines^17,18^, and the DMT1 3’IRE failed to exhibit iron-dependent regulation in reporter assays^18^. Furthermore, *DMT1* transcription is controlled by HIF2α^9,10^, which itself is regulated by IRPs^19^, confounding the study of specific functions of the DMT1 3’IRE^20^. Here, we address the role of this RNA motif using a mouse model with selective disruption of the DMT1 3’IRE.

## Methods

### Animal studies

The *Dmt1* locus was mutagenized through homologous recombination in embryonic stem cells (Supplemental Figure 1). Mice on a mixed 129SvEV/C57BL6 background were housed at the DKFZ under specific-pathogen-free conditions, with unlimited access to standard food and water. All animal work was conducted according to institutional guidelines. Hematological and serum parameters, respectively, were determined using an ABC Vet apparatus (HORIBA ABX SAS, Montpellier, France) and an Olympus-400 analyser (Olympus, Tokyo, Japan). Hepcidin levels were measured using the Hepcidin Murine-Compete™ ELISA kit (Intrinsic LifeSciences, LaJolla, CA).

### RNA and protein analyses

RNA was extracted and used for SYBR Green-based qRT-PCR as described previously^20^. For western blotting, proteins were extracted as described^20^ and analyzed using the antibodies listed in Supplemental Table 2. Tissue staining was performed on paraffin sections using the AEC kit (Vector Lab Inc., Burlingame, CA). Polyribosome analysis was performed as described previously^20^.

### Tissue iron

Tissue non-heme iron levels were measured using the bathophenanthroline method^20^. Iron was stained on paraffin sections using Prussian blue^20^.

## Results and Discussion

We established a mouse line lacking the 5’ stem, the apical loop and part of the 3’ stem of the DMT1 3’IRE (Figure 1A, Supplemental Figure 1). The resulting allele (designated *Dmt1*^IREΔ^) is inherited in Mendelian proportions. Both *Dmt1*^IREΔ/Δ^ males and females are viable and fertile, and display no overt abnormality. We did not observe any major alteration of blood cell parameters (Supplemental Table 1) during postnatal growth (2 weeks of age, during a period of high iron demand), early adulthood (3 months of age) or advanced age (9 months). Hence, while *Dmt1* is essential during perinatal life and critical for erythroid iron acquisition^3-5^, its 3’IRE is not required under standard laboratory conditions and appears to be dispensable for normal hematopoiesis.

**Figure 1:**
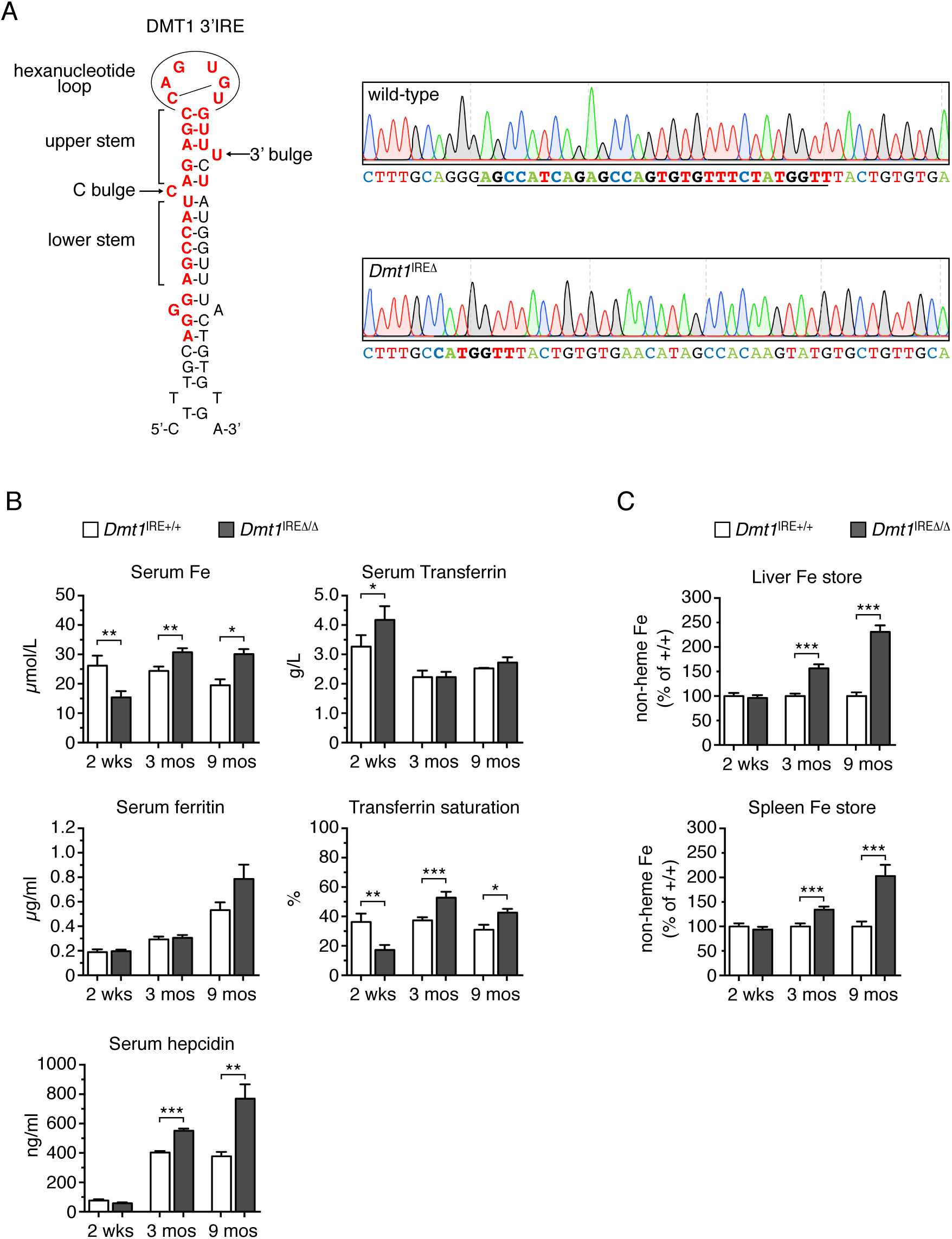
Disruption of the DMT1 3’IRE decreases serum iron levels during early life but leads to hyperferremia in adulthood. (A) Left: Schematic representation of the murine DMT1 IRE stem-loop structure. The bases highlighted in red were deleted by homologous recombination in embryonic stem cells (Supplemental Figure 1). Right: total RNA from wild-type (top) and mutant (bottom) littermates was reverse transcribed and the IRE region of the DMT1 mRNA was PCR amplified (Supplemental Table 2) and sequenced. The electrophoregrams confirm proper mutagenesis of the 3’IRE of the DMT1 mRNA. (B) Serum iron parameters were assessed in male mice at 2 weeks (2 wks), 3 months (3 mos) and 9 months (9 mos) of age (6 to 13 male mice per group). (C) The liver and splenic iron stores were determined in male mice (10 to 22 mice per group) at different stages of life, as in (B). The results are expressed as percentage of wild-type littermates for each range of age. Note that the hepatic and splenic iron stores of 2 week-old mice are low compared to adults (liver: ∼ 100 ±32 mg non-heme iron /g of dry tissue in 2-week old wild-type males versus 279 ±51 and 357 ±62 mg/g at 3 and 9 months of age, respectively; spleen: 227 ±14 mg/g in 2 week-old wild-type males versus 1927 ±119 and 3887 ±744 mg/g at 3 and 9 months of age, respectively). Histograms display averages ± SEM. p: Student’s t-test (*: p<0.05; **: p<0.01; ***: p<0.001).

Interestingly, 2 week-old *Dmt1*^IREΔ/Δ^ male mice display a 40% reduction in serum iron levels, and a decrease in transferrin saturation (Figure 1B). Hepcidin concentration, which is low during postnatal growth, is not affected. Similarly, both hepatic and splenic iron stores are indistinguishable from wild-type animals (Figure 1C). In contrast to suckling pups, 3 month-old *Dmt1*^IREΔ/Δ^ males exhibit high serum iron and transferrin saturation values (Figure 1B) and an increase in tissue iron stores (Figure 1C), accompanied by an augmentation of serum hepcidin (Figure 1B) attributable to stimulation of liver hepcidin via BMP/SMAD signaling (Supplemental Figure 2)^13^. 9 month-old *Dmt1*^IREΔ/Δ^ males display a comparable hyperferremia and a trend towards high serum ferritin (Figure 1B). *Dmt1*^IREΔ/Δ^ female mice show a similar, albeit less pronounced iron phenotype (Supplemental Figure 3) at 2 weeks and 3 months. Enlargement of tissue iron stores in *Dmt1*^IREΔ/Δ^ adults is not associated with aberrantly high expression of known iron import or sequestration genes, nor with marked suppression of molecules involved in iron export (Supplemental Figure 4), suggesting that tissue iron loading is secondary to the elevation of serum iron rather than a consequence of aberrant iron management in liver and spleen cells.

Iron dyshomeostasis in *Dmt1*^IREΔ/Δ^ mice could result from altered intestinal iron absorption^5^. At 2 weeks of age, duodenal non-heme iron levels are unchanged (Figure 2A). However, *Dmt1*^IREΔ/Δ^ pups exhibit a selective downregulation of the DMT1-IRE mRNA isoform, associated with a decrease in total DMT1 protein levels and reduced DMT1 immunostaining at the apical membrane of enterocytes (Figure 2B). This is in agreement with the predicted role of 3’UTR IREs, based on analogy to TFRC, where IRP binding decreases mRNA decay^13,14^. Considering that FPN protein and HEPH mRNA levels are normal (Figure 2C,D), the hypoferremia in *Dmt1*^IREΔ/Δ^ pups could be explained by downregulation of intestinal DMT1 expression.

**Figure 2:**
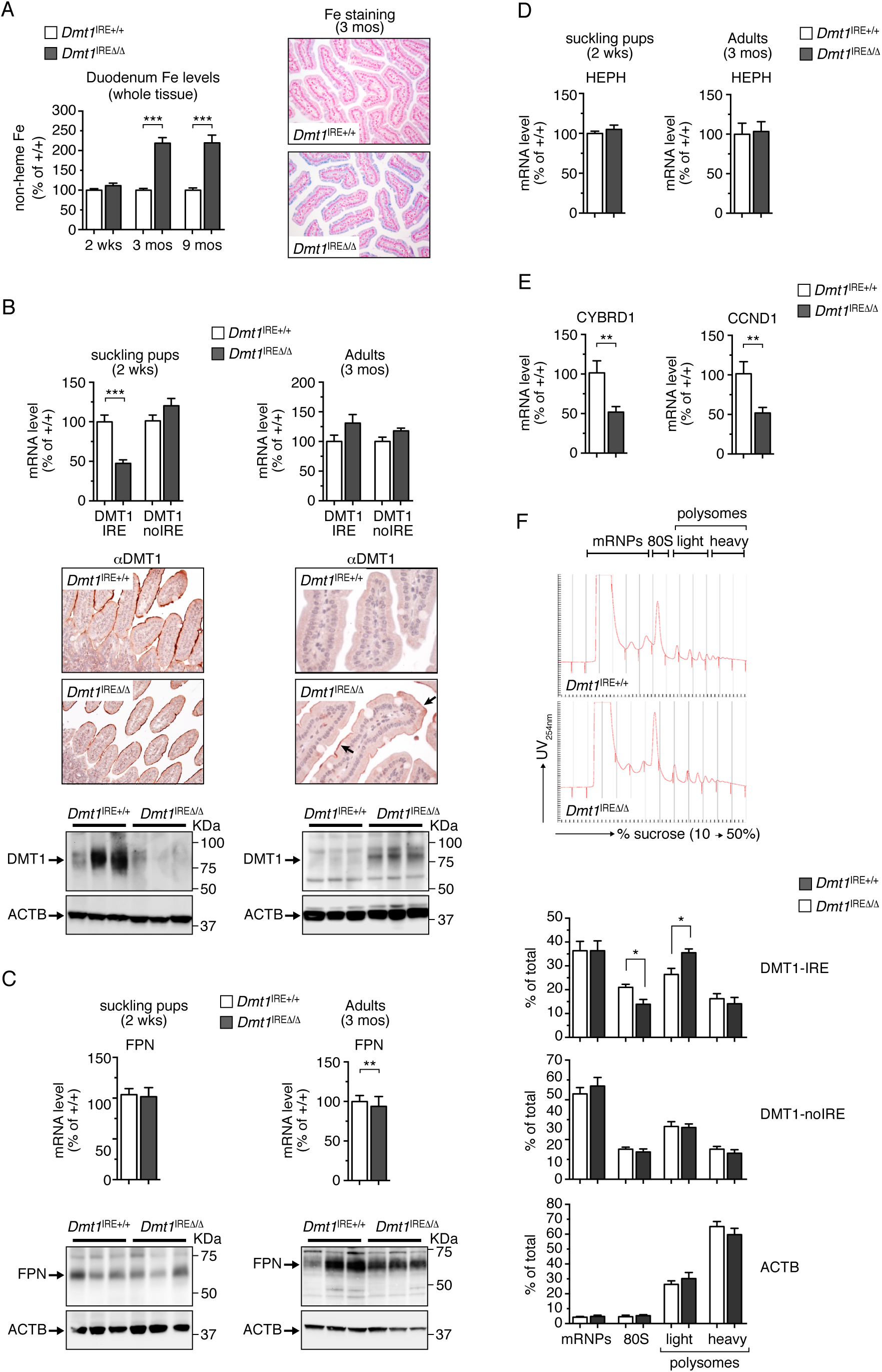
The DMT1 3’IRE exerts divergent effects on intestinal DMT1 expression in postnatal versus adult animals. (A) Left: Duodenal non-heme iron levels were analyzed in male mice at different stages of life as in Figure 1C (9 to 15 mice per group). Right: Perl’s staining in the duodenum of 3 month-old male mice showing iron deposition in mucosal cells (counterstain: nuclear fast red). (B) Top: qPCR analysis of DMT1 mRNA levels in the duodenum of 2 week-(left) versus 3 month-(right) old mice using IRE versus non-IRE-specific primers (Supplemental Table 2). Middle panels: immunostaining of DMT1 in the duodenum (counterstain: hematoxylin); a higher magnification is presented for adult tissues because the DMT1 signal (arrows) is fainter than in suckling pups. Bottom: representative western-blot analysis of DMT1 in whole duodenal extracts. (C) qPCR (top) and western-blot (bottom) analysis of FPN expression in the duodenum of mice at 2 weeks (left) versus 3 months (right) of age. For western blotting (B and C), ACTB was used as loading control. In (A) and (B), pictures were acquired with a DM5000 microscope equipped with a DFC420C camera and a 10X (A), 20X (B, 2 wks) or 40X (B, 3mos) objectives, and were processed using the Leica Application Suite (Leica Biosystems, Wetzlar, Germany). (D) qPCR analysis of HEPH mRNA levels in the duodenum of 2 week-versus 3-month old mice. (E) qPCR analysis of HIF2-target genes in the duodenum of 3 month-old animals. The qPCR data in (B) to (E) display transcript levels as percentage of control (i.e. *Dmt1*^IRE+/+^) after calibration to ACTB (n= 11 to 16 mice per group); similar results were obtained when using GAPDH and/or TUBB5 as standard (not shown). (F) Duodenal scrapings from 3 month-old mice were resolved through linear 20%–50% sucrose density gradients^20^ and fractions were collected with constant UV recording (top). Fractions corresponding to untranslated mRNA (mRNP: messenger ribonucleoprotein), monosomes (80S), as well as light and heavy polysomes were collected. The amount of DMT1 (IRE and non-IRE isoforms) and ACTB mRNAs in those fractions was determined by qPCR. The histograms (bottom) represent mRNA levels across the fractions as a percentage (mean ±SEM, n = 7 to 8) of the sum of all fractions. Histograms display averages ± SEM. p: Student’s t-test (*: p<0.05; **: p<0.01; ***: p<0.001).

During adulthood, *Dmt1*^IREΔ/Δ^ mice exhibit an approximately 2-fold increase in enterocyte iron accumulation (Figure 2A), associated with normal expression of FPN and HEPH (Figure 2C,D). In contrast to its partial suppression in *Dmt1*^IREΔ/Δ^ pups, IRE-containing DMT1 mRNA is expressed at nearly wild-type levels in adult intestine (Figure 2B). It is unlikely that this is due to compensatory stimulation of *Dmt1* transcription by HIF2^9,10^. Indeed, the mRNA levels of *Fpn* and *Ccnd1*, both HIF2-target genes, are not increased (Figure 2C, E). *Cybrd1*, another HIF2-target, appears to be repressed (Figure 2E). Hence, the 3’IRE of DMT1 exerts a positive effect on intestinal DMT1 expression during postnatal growth but not during adult life. This age-dependent effect of the DMT1 3’IRE appears to be tissue specific. While it is also observed in heart, it is not detected in spleen or kidney (Supplemental Figures 4 and 5). Surprisingly, while *Dmt1*^IREΔ/Δ^ adults express nearly wild-type levels of DMT1-IRE mRNA, duodenal DMT1 protein expression is increased (Figure 2B). Although we cannot exclude changes in protein turnover^21^, we speculated that disruption of the DMT1 3’IRE could alter mRNA translation. Supporting this notion, we observed a significant shift of the DMT1-IRE mRNA isoform from monosomes to polysomes (Figure 2F), suggesting that the DMT1 3’IRE partially represses DMT1 mRNA translation in the adult duodenum.

There has been doubt about the functionality of the DMT1 3’IRE. Although it exhibits weaker affinity for IRPs than other IREs^22^, the DMT1 3’IRE does bind IRP1 in the native cellular environment^23^. Our work now demonstrates that the DMT1 3’IRE functions to maintain systemic iron homeostasis, exerting distinct effects at different life stages. It is well recognized that DMT1 regulation in the duodenal mucosa is strongly dependent on HIF2^9,10,24^. In that context, the precise role of the age-dependent switch in the activity of the DMT1 3’IRE remains to be defined. Conceivably, the DMT1 3’IRE may contribute to maintaining baseline homeostasis of immature intestinal absorption early in life. In adulthood, the 3’IRE may help to fine-tune DMT1 expression to avoid excessive iron assimilation.

## Acknowledgements

We thank Sandro Altamura (University of Heidelberg, Germany) for fruitful discussions. We are grateful to the staff of the DKFZ animal facility for their dedicated care of the animals. We thank the “Plateforme de Biochimie” at the “Centre de Recherche sur l’Inflammation” (Paris, France) for their measurement of serum parameters. This work was supported by the Howard Hughes Medical Institute (N.C.A.) and a grant from the Deutsche Forschungsgemeinschaft to B.G. (GA2075/5-1).

## Authorship Contributions

Contribution: designed the research: B.G., N.C.A.; performed experiments: E.T., H.G., S.G., T.B., F.C., M.B., G.P., M.S., B.G.; provided critical reagents: Z.K.; analyzed the data: E.T., S.G., T.B., M.B., Z.K., M.S., N.C.A., B.G.; wrote and reviewed the manuscript: E.T., T.B., Z.K., N.C.A., B.G.

## Conflict-of-interest disclosure

The authors declare no competing financial interests.

