## Supplemental for "Control of Systemic Iron Homeostasis by the 3’ Iron-Responsive Element of Divalent Metal Transporter 1 in Mice"

Supplemental Figure 1: **Targeted mutagenesis of the 3'IRE of the murine *Dmt1* gene.**

Supplemental Figure 2: **Regulation of liver hepcidin in adult mice lacking the 3'IRE of DMT1.**

Supplemental Figure 3: **Iron parameters in female mice.**

Supplemental Figure 4: **Expression of key iron metabolism molecules in the liver and spleen of adult mice**.

Supplemental Figure 5: **DMT1 expression in heart and kidney tissues.**

Supplemental Table 1: **Baseline hematological parameters in *Dmt1*$^{IRE\Delta/\Delta}$ mice are globally normal.**

Supplemental Table 2: **List of oligonucleotides and antibodies used in the study.**

Supplemental References:

Supplemental Figure 1

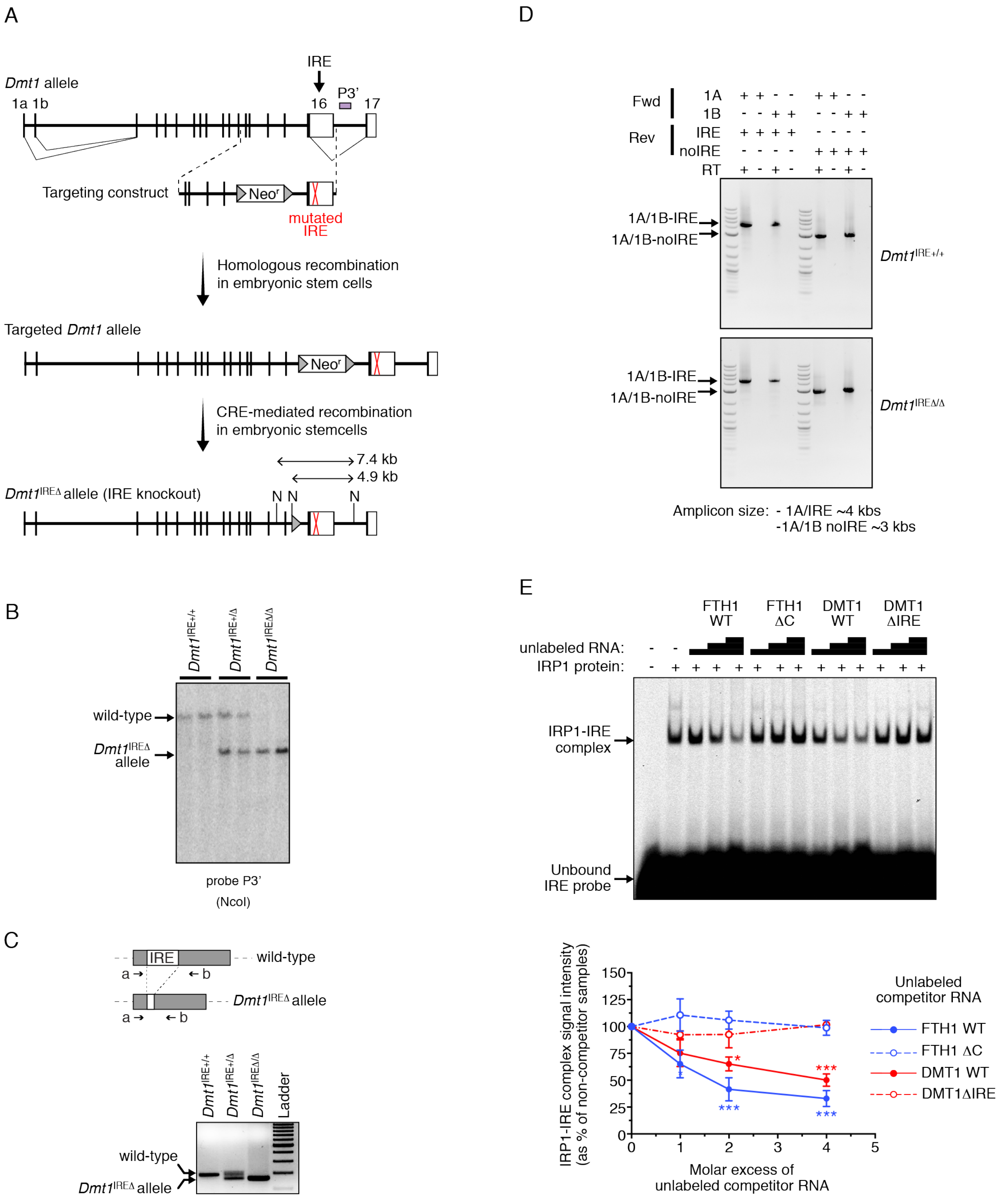


**Supplemental Figure 1: Targeted mutagenesis of the 3'IRE of the murine *Dmt1* gene.**

(A) Schematic representation of the 18 exonic sequences of the *Dmt1* locus and of the strategy used to mutagenise the 3'IRE. The first exon is transcribed from two alternative promoters termed 1A and 1B, respectively. 3' end diversity results from alternative splicing and alternative usage of polyadenylation sites: inclusion of exon 16 gives rise to DMT1 mRNA isoforms bearing the 3'UTR IRE; non-IRE isoforms arise from the joining of a splice donor located upstream of the IRE within exon 16 with the splice acceptor of exon 17. We mutagenised the IRE stem-loop (Figure 1A) using a targeting construct encompassing exons 12 to 16; a cytidine deaminase (CD) neomycin (Neo) resistance cassette (indicated Neo$^r$) flanked with *LoxP* sites (depicted by grey triangles) was inserted into intron 15 and served as a selection marker. The *Dmt1* IRE was replaced by the mutated IRE through homologous recombination in embryonic stem cells. The CD-Neo cassette was subsequently removed via transient expression of CRE recombinase to obtain the *Dmt1*$^{IRE\Delta}$ allele. (B) Southern-blot analysis of *Dmt1*$^{IRE+/+}$, *Dmt1*$^{IRE+/\Delta}$ and *Dmt1*$^{IRE\Delta/\Delta}$ mice with a 3' external probe (P3') after digestion of genomic DNA with NcoI, indicated N in (A). The size of the DNA fragments obtained is indicated. (C) The deletion of the IRE sequence can be detected by genomic PCR using the primers a and b (See Supplemental Table 2) and was used for routine genotyping. (D) To ascertain that mutagenesis of the Dmt1 3'IRE does not lead to overall alterations of transcript synthesis, we reverse transcribed total RNA from intestinal tissue using oligo-dT and PCR amplified all four DMT1 isoforms using primers corresponding to the most distal parts of the RNA. Forward primers (Fwd) are located in exons 1A or 1B, respectively. Reverse primers (Rev) correspond to either exon 16 (IRE) or 17 (noIRE), respectively, and in both cases are located near the polyA site (See Supplemental Table 2). We obtained the amplicons of the expected size (as indicated) from both wild-type (top) and mutant (bottom) mice. Negative control reactions were performed omitting the reverse transcriptase (RT). The boundaries of the PCR products were partially sequenced to confirm the identity of the amplicons. (E) To ascertain that mutagenesis of the DMT1 IRE effectively impairs IRP binding, the last 515 nucleotides of the coding region plus the first 1575 nucleotides of the 3'UTR of the DMT1-IRE mRNA isoform were PCR amplified from *Dmt1*$^{IRE+/+}$ versus *Dmt1*$^{IRE\Delta/\Delta}$ mice (See primers in Supplemental Table 2). The cDNA obtained was sub-cloned and served as template for in vitro synthesis of unlabeled RNA used in a competition electromobility shift assay as described previously.[1] Competitor transcripts corresponding to the the full length FTH1 mRNA (FTH1 WT) versus a mutant version lacking the C residue in the IRE loop (FTH1 ΔC) were used as control to validate the assay. As expected, the unlabeled wild-type FTH1 mRNA competes for the interaction between recombinant purified IRP1 and a fluorescently labeled FTH1 IRE probe, whereas the mutant version does not.[1] Similarly, the wild-type DMT1 RNA (DMT1 WT) competes efficiently, but not the RNA from *Dmt1*$^{IRE\Delta/\Delta}$ mice (DMT1 ΔIRE), confirming efficient disruption of the DMT1 3'IRE. The graph displays the data as average ±SEM relative to the reaction without competitor (p: Student's t-test; *: $p<0.05$; ***: $p<0.001$).

Supplemental Figure 2

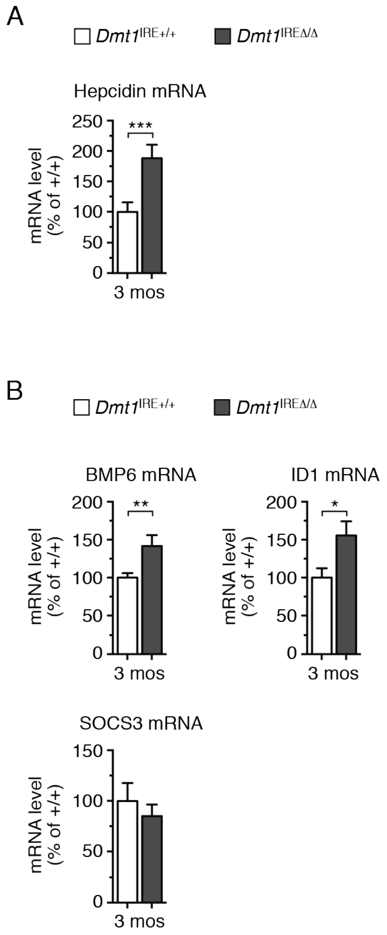


**Supplemental Figure 2: Regulation of liver hepcidin in adult mice lacking the 3'IRE of DMT1.**
(A) qPCR analysis showing an almost 2-fold augmentation of hepcidin mRNA expression in the liver of 3 month-old *Dmt1*$^{IRE\Delta/\Delta}$ male mice compared to wild-type littermates. (B) The upregulation of hepcidin is mirrored by an increase in bone morphogenetic protein 6 (*Bmp6*) and inhibitor of DNA binding 1 (*Id1*) transcript levels, suggesting that hepcidin stimulation in the liver is due to activation of the BMP/SMAD pathway as a consequence of hyperferremia.[2] The expression of SOCS3 remains constant and indicates that JAK-STAT signaling is likely not involved in Hepcidin upregulation.[3] Data are presented as average ±SEM relative to wild-type control after calibration to ACTB (n= 10 to 18 mice per group). p: Student's t-test (*: $p<0.05$; **: $p<0.01$; ***: $p<0.001$).

Supplemental Figure 3

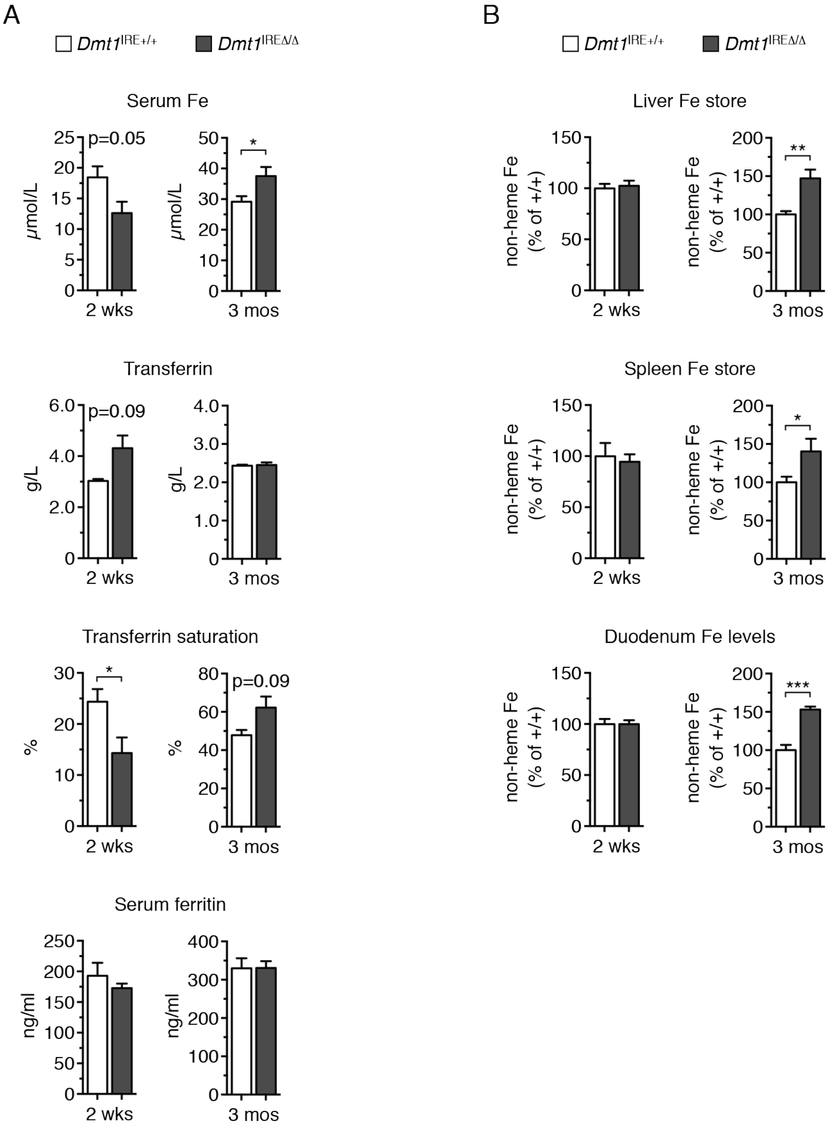


**Supplemental Figure 3: Iron metabolism parameters in female mice**

(A) Serum iron parameters were assessed in female mice during postnatal growth (2 weeks of age) or during early adulthood  (3 months of age), respectively (8 o 13 mice per group). Similar to male mice (Figure 1C), *Dmt1*$^{IRE\Delta/\Delta}$ female mice display a tendency towards decreased serum iron concentration and transferrin saturation during the suckling period (2 wks). On the opposite, they tend to have higher serum and transferrin saturation values during adulthood. Furthermore, the increase in serum iron levels is accompanied by a modest but significant augmentation of the liver and spleen iron stores (B) (10 to 21 mice pr group), as observed in males (Figure 1D). Histograms display averages ± SEM. p: Student's t-test (*: $p<0.05$; **: $p<0.01$; ***: $p<0.001$).

Supplemental Figure 4

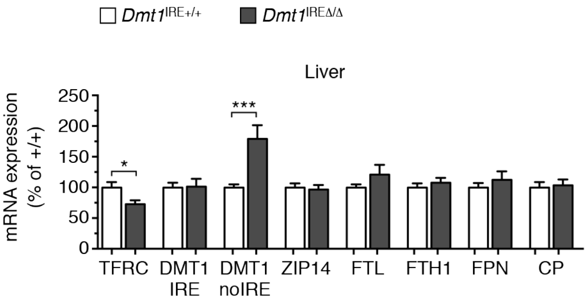


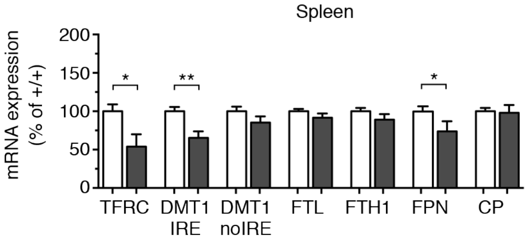


**Supplemental Figure 4: Expression of key iron metabolism molecules in the liver and spleen of adult mice.**

qPCR analysis of iron metabolism molecules in liver (top) and spleen (bottom) tissues of *Dmt1*$^{IRE\Delta/\Delta}$ versus *Dmt1*$^{IRE+/+}$ male mice at 3 months of age. TFRC mRNA levels are reduced in liver, possibly as a response to hepatic iron loading. DMT1-IRE mRNA expression is unaltered, but DMT1-noIRE mRNA levels are increased. Since DMT1-noIRE contributes to iron acquisition downstream of TFRC[4] and TFRC is downregulated in *Dmt1*$^{IRE\Delta/\Delta}$ mice, hepatic iron accumulation unlikely results from stimulation of transferrin-bound iron uptake. ZIP14 (a.k.a. SLC39A14), which mediates the uptake of non-Tf bound iron in liver cells[5], is unchanged. The mRNA levels of the ferritin-L (FTL) and -H (FTH1) iron sequestration molecules are comparable between mutant and wildtype. FPN and the ferroxidase ceruloplasmin (CP), both required for cellular iron export[6,7], are unaltered. In spleen, iron accumulation cannot be explained by increased expression of iron import molecules, as both TFRC and DMT1-IRE are downregulated. We observed a mild reduction (~25%) of FPN mRNA levels, which unlikely suffices to cause iron retention in splenic macrophages since 3 month-old mice lacking one *Fpn* allele display normal iron homeostasis parameters[8]. The FTL and FTH1 mRNAs are unaffected. Overall these data suggest that enlargement of tissue iron stores in adult *Dmt1*$^{IRE\Delta/\Delta}$ animals does not result from iron mismanagement in liver and spleen cells and is rather secondary to the increase in serum iron levels. Data are presented as average ±SEM relative to wild-type control after calibration to ACTB (n= 7 to 17 mice per group). p: Student's t-test (*: $p<0.05$; **: $p<0.01$; ***: $p<0.001$).

Supplemental Figure 5

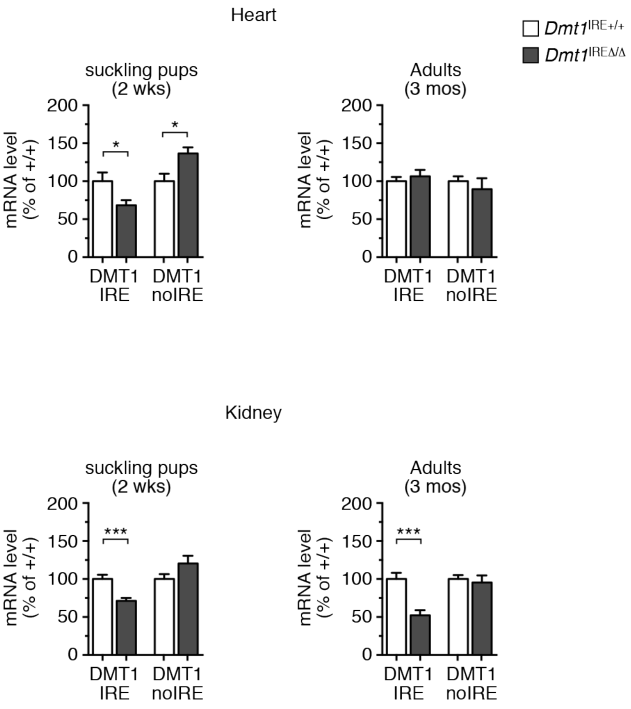


**Suplemental Figure 5: DMT1 expression in heart and kidney tissues during postnatal life and adulthood.** qPCR analysis of DMT1 3' mRNA variants in the heart (top) and kidney (bottom) of *Dmt1*$^{IRE\Delta/\Delta}$ versus *Dmt1*$^{IRE+/+}$ male mice at 2 weeks or at 3 months of age, respectively. In heart, we observe a mild but significant decrease in DMT1-IRE mRNA expression in 2 week-old *Dmt1*$^{IRE\Delta/\Delta}$ mice, associated with an opposite increase in the DMT1 non-IRE mRNA levels. Similar to the duodenum, DMT1 mRNA expression is unaltered when mice reach adulthood. In kidney, DMT1-IRE mRNA expression is reduced both during postnatal growth and early adulthood, suggesting that the activity of 3'IRE of DMT1 is not only age but also tissue dependent. Data are presented as average ±SEM relative to wild-type control after calibration to ACTB (heart: 5 to 6 mice per group; kidney: 9 to 10 mice per group). p: Student's t-test (*: $p<0.05$; ***: $p<0.001$).

**Supplemental Table 1: Baseline hematological parameters in *Dmt1*$^{IRE\Delta/\Delta}$ mice are globally normal.**

| | 2 wks | | 3 mos | | 9 mos | |
|---|---|---|---|---|---|---|
| | ***Dmt1***$^{IRE+/+}$ | ***Dmt1***$^{IRE\Delta/\Delta}$ | ***Dmt1***$^{IRE+/+}$ | ***Dmt1***$^{IRE\Delta/\Delta}$ | ***Dmt1***$^{IRE+/+}$ | ***Dmt1***$^{IRE\Delta/\Delta}$ |
| *Males* | (n=8) | (n=18) | (n=11) | (n=21) | (n=16) | (n=12) |
| WBC ($10^9$/L) | 3.7 ±0.5 | 3.5 ±0.2 | 4.5 ±0.3 | 4.4 ±0.3 | 3.2 ±0.2 | 3.8 ±0.3 |
| RBC ($10^{12}$/L) | 5.5 ±0.3 | 6.1 ±0.1 ** | 8.7 ±0.2 | 8.4 ±0.1 | 8.7 ±0.1 | 8.9 ±0.1 |
| HGB (g/L) | 10.6 ±0.2 | 11.2 ±0.2 | 15.2 ±0.3 | 15.2 ±0.1 | 14.8 ±0.2 | 15.2 ±0.2 |
| HCT (L/L) | 32.7±0.9 | 35.2 ±0.6 * | 46.6 ±0.8 | 45.8 ±0.4 | 46.3 ±0.5 | 47.4 ±0.7 |
| MCV (fL) | 59.1 ±0.9 | 57.6 ±0.5 | 53.8 ±0.7 | 54.4 ±0.4 | 53.4 ±0.5 | 53.1 ±0.7 |
| MCHC (pg/L) | 32.5 ±0.4 | 31.7 ±0.3 | 32.5 ±0.3 | 33.2 ±0.3 | 31.9 ±0.3 | 32.1 ±0.4 |
| PLT ($10^9$/L) | 862 ±131 | 1026 ±41 | 1142 ±43 | 1127 ±53 | 1108 ±44 | 1126 ±48 |
| *Females* | (n=11) | (n=11) | (n=13) | (n=11) | (n=9) | (n=10) |
| WBC ($10^9$/L) | 4.0 ±0.3 | 3.9 ±0.2 | 4.5 ±0.2 | 4.5 ±0.5 | 4.5 ±0.5 | 4.6 ±0.4 |
| RBC ($10^{12}$/L) | 5.7 ±0.2 | 5.6 ±0.1 | 8.6 ±0.1 | 8.7 ±0.1 | 8.5 ±0.3 | 9.0 ±0.2 |
| HGB (g/L) | 11.0 ±0.3 | 10.7 ±0.2 | 15.3 ±0.3 | 15.8 ±0.2 | 14.3 ±0.3 | 14.8 ±0.3 |
| HCT (L/L) | 34.9 ±1.0 | 33.6 ±0.8 | 45.2 ±0.6 | 46.3 ±0.5 | 44.2 ±0.9 | 46.0 ±1.0 |
| MCV (fL) | 60.9 ±0.9 | 60.5 ±1.1 | 52.2 ±0.5 | 52.8 ±0.5 | 52.2 ±1.2 | 51.2 ±0.7 |
| MCHC (pg/L) | 31.7 ±0.4 | 31.8 ±0.3 | 33.9 ±0.5 | 34.2 ±0.4 | 32.3 ±0.3 | 32.3 ±0.2 |
| PLT ($10^9$/L) | 1001 ±30 | 1119 ±33 * | 956 ±54 | 1007 ±76 | 1317 ±38 | 1401 ±32 |

Hematological parameters were assessed in both male (top) and female (bottom) mice at 2 weeks of age, during early adulthood (3 months of age) and during aging (9 months of age), using EDTA blood. *Dmt1*$^{IRE\Delta/\Delta}$ mice display globally unchanged hematological parameters, with only a very mild and transient increase in red blood cell counts and hematocrit values in 2 week-old male pups. This shows that the 3'IRE of DMT1 is largely dispensable for normal erythropoiesis under standard laboratory conditions, and is compatible with the notion that erythroid cells rely on the nonIRE isoform of DMT1 for the uptake of transferrin-bound iron. The fact that the mice do not display signs of iron deficiency anemia in spite of the reduction in serum iron levels might be an indication that the hypoferremia is only transient and/or compensated for by the erythropoietic system.

WBC: white blood cells; RBC: red blood cells; HGB: hemoglobin; HCT: hematocrit; MCV: mean corpuscular volume; MCHC: mean corpuscular hemoglobin concentration; PLT: platelets. Results are presented as means ± standard error. Sample size (n) is indicated.

* $p<0.05$; ** $p<0.01$; unpaired, two-tailed t-test.

### Supplemental Table 2: Primers and antibodies used in the study

| **Oligonucleotides** | | | | |
|---|---|---|---|---|
| *Gene name* | *primer name* | *sequence (5' to 3')* | *purpose* | *note* |
| *Dmt1 (Slc11a2)* | forward<br>reverse | TCCTTGGCACTTGCATACTCAC<br>AGGTAACACGAAAGGCTAAGGTG | Southern blotting | generation of probe P3' in Supplemental Figure 1B |
| | exon 16 primer a<br>exon 16 primer b | GGTAAGCATCTCGAAAGTCC<br>CCCAACTAACAGTTGAGTCC | genomic PCR | routine genotyping in Supplemental Figure 1C |
| | exon 1A forward<br>exon 1B forward<br>exon 16 reverse<br>exon 17 reverse | AAGCCAAACCAGTCCTGCACC<br>AGGGCAGGAGGTTGACTGG<br>GTTCACTAAAGCCTCCTTGAAGC<br>TCTGGTTGGGATTAAAGCAAAAACC | RT-PCR | Analysis of DMT1 mRNA isoforms in Supplemental Figure 1D |
| | exon 12 forward<br>exon 16 reverse | ACCTATTCTGGCCAGTTTGTCATG<br>GGGGACAACTATTTCAGGTC | Cloning | DNA template for in vitro transcription of competitor RNA in Supplemental Figure 1E |
| | qPCR IRE forward<br>qPCR IRE reverse | ATGTTGCCACCGCTGGTATC<br>AGCTAGGCCATGTGGCACTCT | qPCR | analysis of DMT1 IRE mRNA isoforms (Figure 2 + Supplemental Figure 5) |
| | qPCR noIRE forward<br>qPCR noIRE reverse | GCGGTCAGTCCCAGGCGGTACG<br>GTGGTGGCTGCAGTGGTTAGCG | qPCR | analysis of DMT1 noIRE mRNA isoforms (Figure 2 + Supplemental Figure 5) |
| *Fpn (Slc40a1)* | forward<br>reverse | GGGTGGATAAGAATGCCAGACTT<br>GTCAGGAGCTCATTCTTGTGTAGGA | qPCR | Analysis of iron export molecules in Figure 2C |
| *Heph* | forward<br>reverse | TCTATACATGCCCATTGGAGTTCT<br>TGGGATGTTCCACTGGTAAGT | qPCR | Analysis of iron export molecules in Figure 2C |
| *Cybrd1* | forward<br>reverse | GAAAAGCTGTTCTTTGTCCTGAAAC<br>GCCCAGCGTATTTGTAAAAACAC | qPCR | Analysis of HIF2 targets in Figure 2D |
| *Ccnd1* | forward<br>reverse | CATCCATGCGGAAAATCG<br>GCGGGAAGACCTCCTCTT | qPCR | Analysis of HIF2 targets in Figure 2D |
| *Socs3* | forward<br>reverse | ATTTCGCTTCGGGACTAGC<br>AACTTGCTGTGGGTGACCAT | qPCR | Analysis of hepcidin in Supplemental Figure 2 |
| *Bmp6* | forward<br>reverse | ACTGACTAGCGCGCAGGA<br>TGTGGGGAGAACTCCTTGTC | qPCR | Analysis of hepcidin in Supplemental Figure 2 |
| *Id1* | forward<br>reverse | GGCGAGATCAGTGCCTTGG<br>CTCCTGAAGGGCTGGAGTC | qPCR | Analysis of hepcidin in Supplemental Figure 2 |
| *Actb* | forward<br>reverse | GGCCAGGATGGAGCCACCGATC<br>CAGCCATTGCTGACAGGATGCA | qPCR | qPCR standard |
| *Hepcidin (Hamp)* | forward<br>reverse | CCTATCTCCATCAACAGAT<br>TGCAACAGATACCACACTG | qPCR | Analysis of hepcidin in Supplemental Figure 2 |
| *Tfrc* | forward<br>reverse | CCCATGACGTTGAATTGAACCT<br>GTAGTCTCCACGAGCGGAATA | qPCR | Analysis of iron molecules in Supplemental Figure 4 |
| *Zip14 (Slc39a14)* | forward<br>reverse | TGGAACCCTCTACTCCAACG<br>CTGAGGGTTGAAGCCAAAAG | qPCR | Analysis of iron molecules in Supplemental Figure 4 |
| *Cp* | forward<br>reverse | GAAAGGCAGCTTACTTGCTGA<br>TCAAACACTGTGGGAAACAAGT | qPCR | Analysis of iron molecules in Supplemental Figure 4 |
| *Fth1* | forward<br>reverse | TGGAACTGCACAAACTGGCTACT<br>ATGGATTTCACCTGTTCACTCAGATAA | qPCR | Analysis of iron molecules in Supplemental Figure 4 |
| *Ftl* | forward<br>reverse | CGTGGATCTGTGTCTTGCTTCA<br>GCGAAGAGACGGTGCAGACT | qPCR | Analysis of iron molecules in Supplemental Figure 4 |
| **Antibodies** | | | | |
| *protein (host)* | *supplier* | *concentration* | *purpose* | *note* |
| DMT1 (rabbit) | self generated against antigen: MVLDPEEKIPDDGAS GDHGDSC | 1/500 | western blotting | reacts with all DMT1 protein isoforms |
| | | 1/400 | immuno-staining | |
| FPN (rabbit) | alpha-diagnostics | 1/300 | western blotting | affinity purified antibody |
| ACTB (mouse) | Sigma-Aldrich | 1/5000 | western blotting | clone |
